# Non-hierarchical, RhlR-regulated acyl-homoserine lactone quorum sensing in a cystic fibrosis isolate of *Pseudomonas aeruginosa*

**DOI:** 10.1101/755314

**Authors:** Renae L. Cruz, Kyle L. Asfahl, Sara Van den Bossche, Tom Coenye, Aurélie Crabbé, Ajai A. Dandekar

## Abstract

The opportunistic pathogen *Pseudomonas aeruginosa* is a leading cause of airway infection in cystic fibrosis (CF) patients. *P. aeruginosa* employs several hierarchically arranged and interconnected quorum sensing (QS) regulatory circuits to produce a battery of virulence factors such as elastase, phenazines, and rhamnolipids. The QS transcription factor LasR sits atop this hierarchy, and activates the transcription of dozens of genes, including that encoding the QS regulator RhlR. Paradoxically, inactivating *lasR* mutations are frequently observed in isolates from CF patients with chronic *P. aeruginosa* infections. In contrast, mutations in *rhlR* are rare. We have recently shown that in CF isolates, the QS circuitry is often “rewired” such that RhlR acts in a LasR-independent manner. To begin understanding how QS activity differs in this “rewired” background, we characterized QS activation and RhlR-regulated gene expression in *P. aeruginosa* E90, a LasR-null, RhlR-active chronic infection isolate. In this isolate, RhlR activates the expression of 53 genes in response to increasing cell density. The genes regulated by RhlR include several that encode virulence factors. Some, but not all, of these genes are present in the QS regulon described in the well-studied laboratory strain PAO1. We also demonstrate that E90 produces virulence factors at similar concentrations to that of PAO1. Unlike PAO1, cytotoxicity by E90 in a three-dimensional lung epithelium cell model is also RhlR-regulated. These data illuminate a “rewired” LasR-independent RhlR regulon in chronic infection isolates and suggest that RhlR may be a target for therapeutic development in chronic infections.

**AUTHOR SUMMARY:** *Pseudomonas aeruginosa* is a prominent cystic fibrosis (CF) pathogen that uses quorum sensing (QS) to regulate virulence. In laboratory strains, the key QS regulator is LasR. Some isolates from patients with chronic CF infections appear to use an alternate QS circuitry in which another transcriptional regulator, RhlR, mediates QS. We show that a LasR-null CF clinical isolate engages in QS through RhlR and remains capable of inducing cell death in an *in vivo*-like lung epithelium cell model. Our findings support the notion that LasR-null clinical isolates can engage in RhlR QS and highlight the centrality of RhlR gene regulation in chronic *P. aeruginosa* infections.

## INTRODUCTION

Many species of bacteria are able to sense and communicate with each other via quorum sensing (QS), a cell-density dependent gene regulation mechanism[1]. In *Proteobacteria*, acyl-homoserine lactones (HSL) are used as QS signals. Commonly, signals are produced by acyl-HSL synthases of the *luxI* family and are recognized by their cognate receptors, transcription factors of the *luxR* family[2].

*Pseudomonas aeruginosa*, a leading cause of airway infection in cystic fibrosis (CF) patients, uses QS to regulate the production of a wide array of virulence factors including phenazines, rhamnolipids, and hydrogen cyanide[3]. *P. aeruginosa* possesses two complete LuxI/LuxR QS regulatory circuits: LasI/LasR and RhlI/RhlR[4,5]. The signal synthase LasI produces the signal *N*-3-oxo-dodecanoyl-homoserine lactone (3OC12-HSL). Above a certain concentration, 3OC12-HSL binds to and facilitates the dimerization of LasR[6]. The LasR homodimer functions as a transcriptional activator promoting the expression of hundreds of genes including *rhlR* and *rhlI*, thereby linking the two acyl-homoserine lactone (AHL) QS regulatory circuits[4,5]. Similarly, RhlI produces the signal *N*-butanoyl-homoserine lactone (C4-HSL), which binds to RhlR, initiating transcription of an additional set of target genes that overlap somewhat with the LasR regulon[1,7]. There is a third, non-AHL QS circuit in *P. aeruginosa* that involves a quinolone signal *(Pseudomonas* quinolone signal; PQS), which activates the transcription factor PqsR[8]. PqsR and RhlR co-regulate the production of some extracellular products[9].

In laboratory strains of *P. aeruginosa*, deletion or deleterious mutation of *lasR* results in attenuated virulence in various animal models of infection[10,11]. Despite the importance of LasR in regulating virulence, several studies have shown that *lasR* mutations are commonly observed in isolates collected from the lungs of chronically infected CF patients[12–14]. In some patients, the frequency of isolates with a mutant *lasR* has been reported to be greater than 50%[13,15]. These findings led to the notion that QS is not essential during chronic stages of infection, dampening enthusiasm for QS inhibitors as potential therapeutics. Contrary to this idea, we and others have shown that many LasR-null *P. aeruginosa* chronic infection isolates remain capable of engaging in QS activity through the RhlI/RhlR circuit[16–18]. CF strains appear to “rewire” their QS circuitry so that RhlR is the key transcription factor.

We are interested in the regulatory remodeling of QS that occurs in isolates of *P. aeruginosa* from chronic infections, including those in CF. To begin to understand how RhlR-mediated QS in clinical isolates might be different from that of laboratory strains[15,16,18,19], we studied a CF isolate called E90[20], which contains a single base-pair deletion in *lasR* at base 170, and uses RhlR to mediate QS. E90 produces QS-regulated virulence factors at levels comparable to that of PAO1. We used RNA-seq to analyze the RhlR regulon of this isolate by comparing its transcriptome with that of an isogenic RhlR deletion mutant. We determined that the E90 RhlR regulon consists of over 83 genes including those that encode virulence factors. Using a three-dimensional tissue culture model, we also observed that E90 induces cell death in a RhlR-dependent manner. Together our data provide a more complete picture of the “rewiring” of QS that can take place in CF-adapted *P. aeruginosa*, while also providing a basis for understanding the gene targets of RhlR without the confounding effects of the QS hierarchy.

## RESULTS

### RhlR and C4-HSL-dependent QS activity is conserved in LasR-null isolate E90

We identified isolate E90 from a phenotypic survey of chronic infection isolates collected in the Early *Pseudomonas* Infection Control (EPIC) Observational Study [16]. This isolate, an apparent LasR mutant, still engaged in activities that are putatively QS-regulated such as rhamnolipid, exoprotease, and phenazine production. The *lasR* gene of E90 features a 1 bp deletion at nucleotide position 170, a frameshift mutation which results in a premature stop codon (at residue 114). To confirm that this single nucleotide polymorphism encodes a nonfunctional LasR polypeptide, we transformed the strain with a LasR-specific reporter plasmid consisting of *gfp* fused to the promoter region of *lasI*, which encodes the signal synthase and is strongly activated by LasR [21]. GFP fluorescence in E90 transformed with this reporter plasmid was nil and mirrored that of a PAO1Δ*lasR* mutant (Fig. 1A). As a complementary approach, we measured the concentration of 3OC12-HSL produced by E90 using a bioassay. We found that E90 after overnight growth produced very low amounts of 3OC12-HSL (40 nM) compared to PAO1 (1.5 μM) and in contrast to PAO1Δ*lasR* for which no 3OC12-HSL was detected (Fig. 1B). The apparent differences in E90 LasR activity reported by the transcriptional reporter assay and the bioassay is likely explained by the greater sensitivity of the latter method. E90 produced approximately 8.3 μM C4-HSL after overnight growth, comparable to what we measured for PAO1 (9.8 μM). Altogether, these data confirmed that E90 encodes a non-functional LasR, and suggested that QS in this isolate is regulated by either RhlR, PqsR, or both transcription factors.

**Figure 1.**
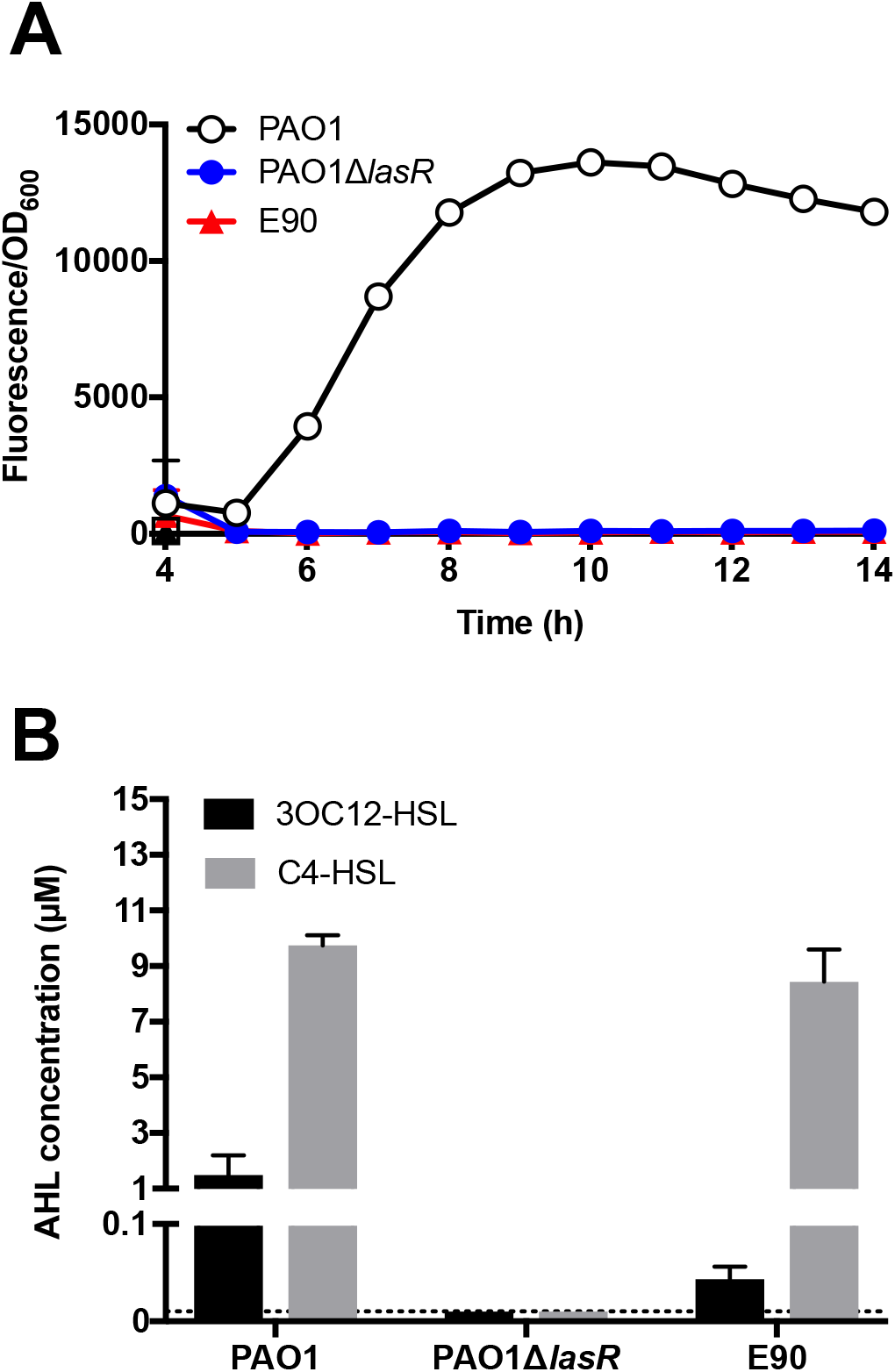
LasR activity is absent in E90, a cystic fibrosis-adapted chronic infection isolate. A) *plasI-gfp* reporter activity over time (Fluorescence/OD_600_). PAO1, open circles; PAO1Δ*lasR*, blue circles; E90, red triangles. Data from the first three hours are excluded from analysis because cell density measurements were below the limit of detection. B) AHL signal concentrations. Black bars, 3OC12-HSL; grey bars, C4-HSL. The dashed line indicates the limit of detection for the 3OC12 and C4-HSL bioassay (10 nM in each case). Both the PAO1Δ*lasR* mutant and E90 produce concentrations of 3OC12-HSL and C4-HSL that are significant from PAO1 (p-value < 0.05 by t-test). For (A) and (B), means and standard deviation of biological replicates are shown (*n*=3). In some cases, error bars are too small to be seen.

To determine if RhlR QS activity in E90 was AHL-dependent, we examined the expression of several well-studied quorum-regulated genes in the presence or absence of AiiA lactonase, an enzyme that degrades AHL signals[22]. Using qRT-PCR, we observed that expression of *lasB* and *rhlA* were increased in the presence of AHLs (Fig. 2). These genes, which encode the exoprotease elastase and a rhamnosyltransferase involved in rhamnolipid production, were identified as QS-regulated in PAO1 [3,23]. *rhlI*, which encodes the C4-HSL synthase, was also AHL-regulated (Fig. 2). Because we had determined that little 3OC12-HSL is produced by E90 (Fig. 1), we reasoned that expression of QS-regulated genes likely was dependent on C4-HSL.

**Figure 2.**
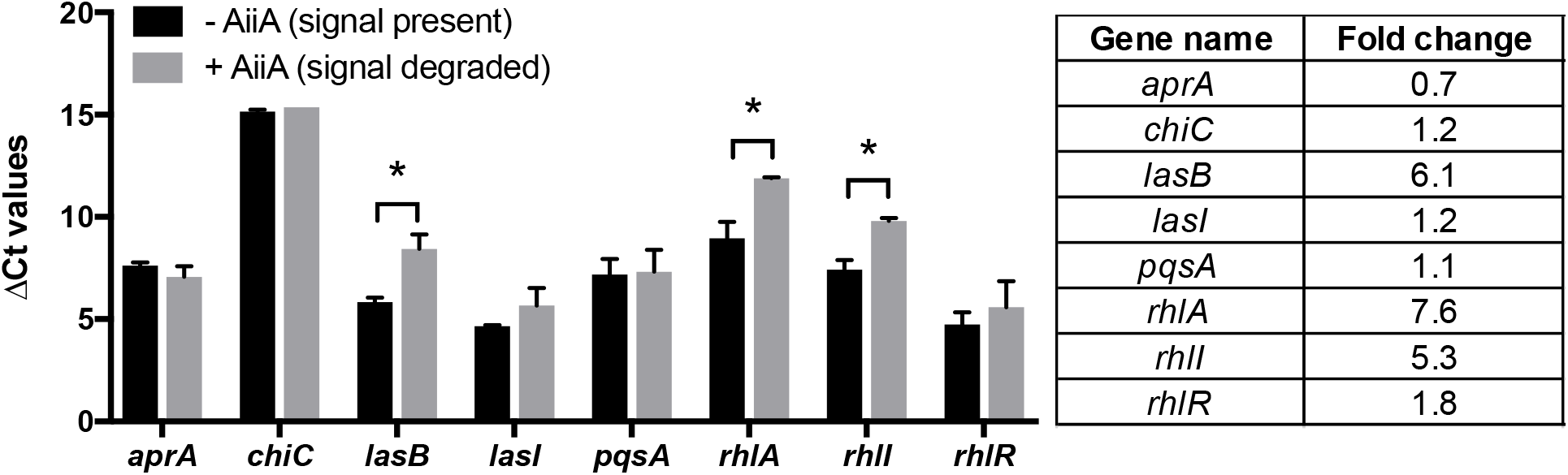
In isolate E90, expression of several canonical QS-regulated genes is AHL-dependent. The following target genes were measured in the presence or absence of AiiA lactonase using qRT-PCR: *lasI*, 3OC12-HSL signal synthase; *lasB*, elastase; *rhlI*, C4-HSL signal synthase; *rhlR*, RhlR, *pqsA*, coenzyme A ligase involved in *Pseudomonas* Quinolone Signal synthesis; *chiC*, chitinase; *aprA;* alkaline metalloprotease. The differences in threshold cycle (Δ Ct) are measured relative to the housekeeping gene *rplU*. Fold changes in gene expression (table on right) are reported relative to cultures incubated with AiiA lactonase. Asterisks (*) indicate statistical significance (p < 0.05 by *t* test). Error bars represent the standard deviation for results of three independent experiments.

Previous work has shown that RhlR activity can be uncoupled from LasR regulation in LasR-null backgrounds [24–26]. Given that C4-HSL production is robust in E90 (Fig. 1B), we queried if this strain similarly engaged in RhlR-dependent QS activity. To address this question, we engineered a *rhlR* deletion in the E90 background to observe its effect on quorum-regulated phenotypes. We found that the E90Δ*rhlR* deletion mutant displayed nil *rhlA* promoter activity and produced little to no exoprotease and pyocyanin, consistent with the idea that RhlR regulates QS activity in E90 (Fig. 3). As a whole, our results showed that the LasR-null isolate E90 retains QS activity in a RhlR- and C4-HSL-dependent manner, and suggested that regulation by RhlR in this strain parallels that of LasR in PAO1. Because RhlR-dependent QS regulation appears to be common in CF isolates[16–18], we reasoned that a study of the genes regulated by RhlR in this background would give insight into which QS-regulated gene products might be important in chronic CF infections. Furthermore, because *rhlR* is not regulated by LasR in E90 (and other clinical isolates), a study of the E90 QS transcriptome has the potential to disentangle genes that are regulated solely by RhlR from those that require both LasR and RhlR.

**Figure 3.**
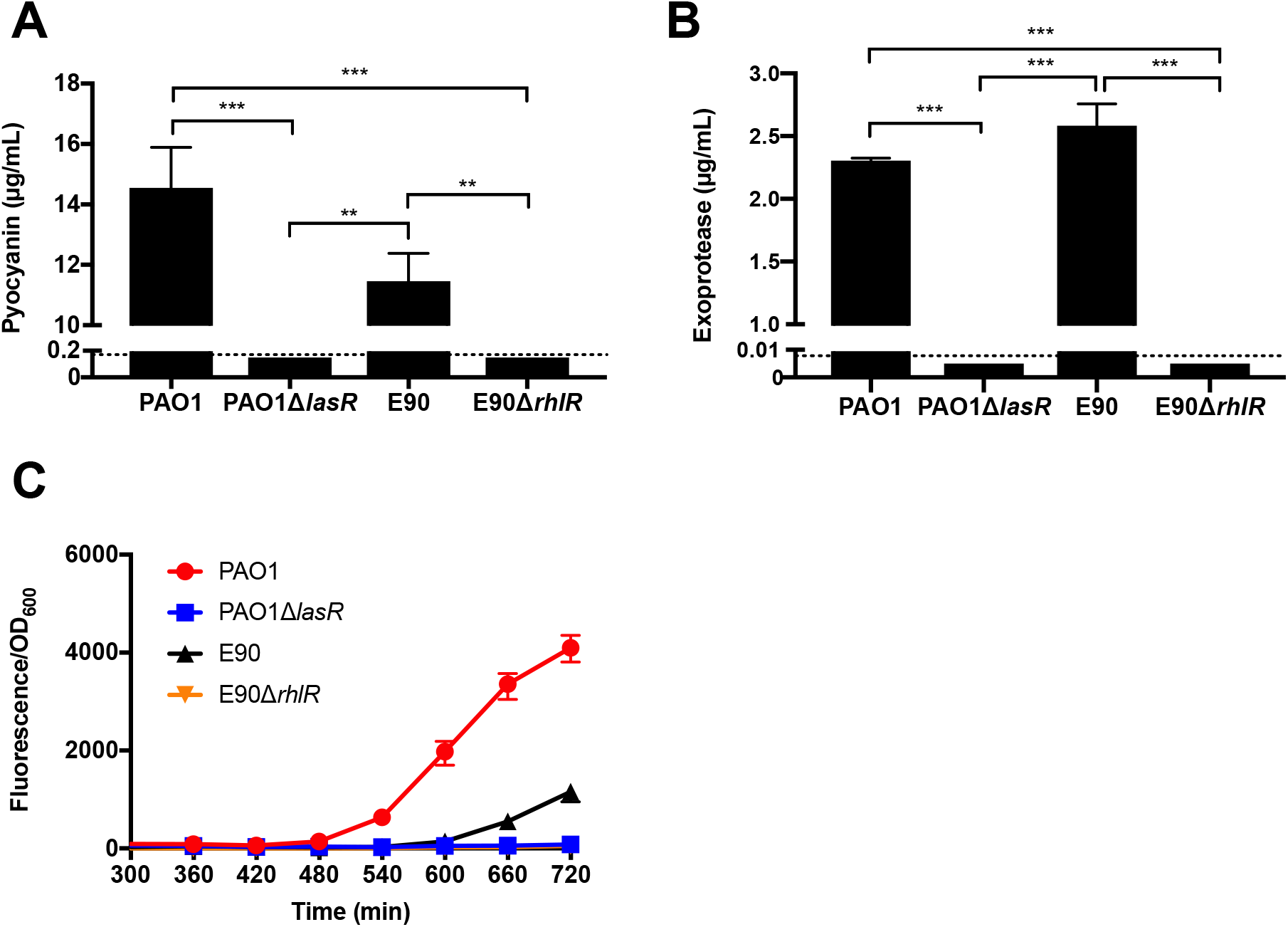
RhlR regulates QS in E90. Production of (A) pyocyanin or (B) protease in either PAO1, E90, PAO1Δ*lasR* or E90Δ*rhlR*. The dashed lines indicate the detection limits for the pyocyanin and protease assays, which are 0.2 μg/mL and 0.008 μg/mL, respectively. C) *rhlA-gfp* reporter activity over time. Data from the first five hours are excluded because cell density measurements were below the limit of detection of the plate reader. Error bars represent the standard deviation for results of three independent experiments. In some cases, error bars are too small to be seen. Both the PAO1Δ*lasR* mutant and E90 produce concentrations of pyocyanin or exoprotease that are significantly different from PAO1 (*, p < 0.05; **, p < 0.01; ***, p < 0.001 by t-test).

### Identification of the RhlR regulon of E90

To determine which genes are regulated by RhlR, we performed an RNA-seq-based differential gene (DE) expression analysis comparing RNA collected from cultures of the parent strain E90 to the isogenic RhlR deletion mutant. First, we sought to generate a *de* novo-assembled genome for E90 to use as an RNA-seq mapping reference which would account for the potential genomic differences between E90 and reference strains of *P. aeruginosa*. Using a hybrid approach combining both short- and long-read high-throughput sequencing, we were able to assemble the genome of E90 into a single circular contig of approximately 6.8 Mb that harbors 6650 annotated features (Figure 4; 6650 features total, 6503 protein coding sequences). In addition to being roughly 550 kb larger than the published sequence of laboratory strain PAO1 [27], the genome of E90 includes 862 features with no homology to PAO1. Also present is a 4.4 Mb inversion relative to PAO1, which includes an internal reorder of roughly 250 kb. The inversion appears to be the result of a recombination event between two roughly 5 kb repeat regions that do not have homology to PAO1, but flank the *rrnA/rrnB* region previously implicated in restructuring of the *P. aeruginosa* genome [27]. A brief search of the E90 genome for *P. aeruginosa* genes previously reported to be under purifying selection in CF isolates revealed a nonsynonymous mutation in the gene coding for the probable oxidoreductase MexS (locus PAE90_2949/PA2491; nonsynonymous SNP), as well as the resistance-nodulation-division multidrug efflux membrane fusion protein precursor MexA (locus PAE90_0464/PA0425; 33bp deletion) [13].

**Figure 4.**
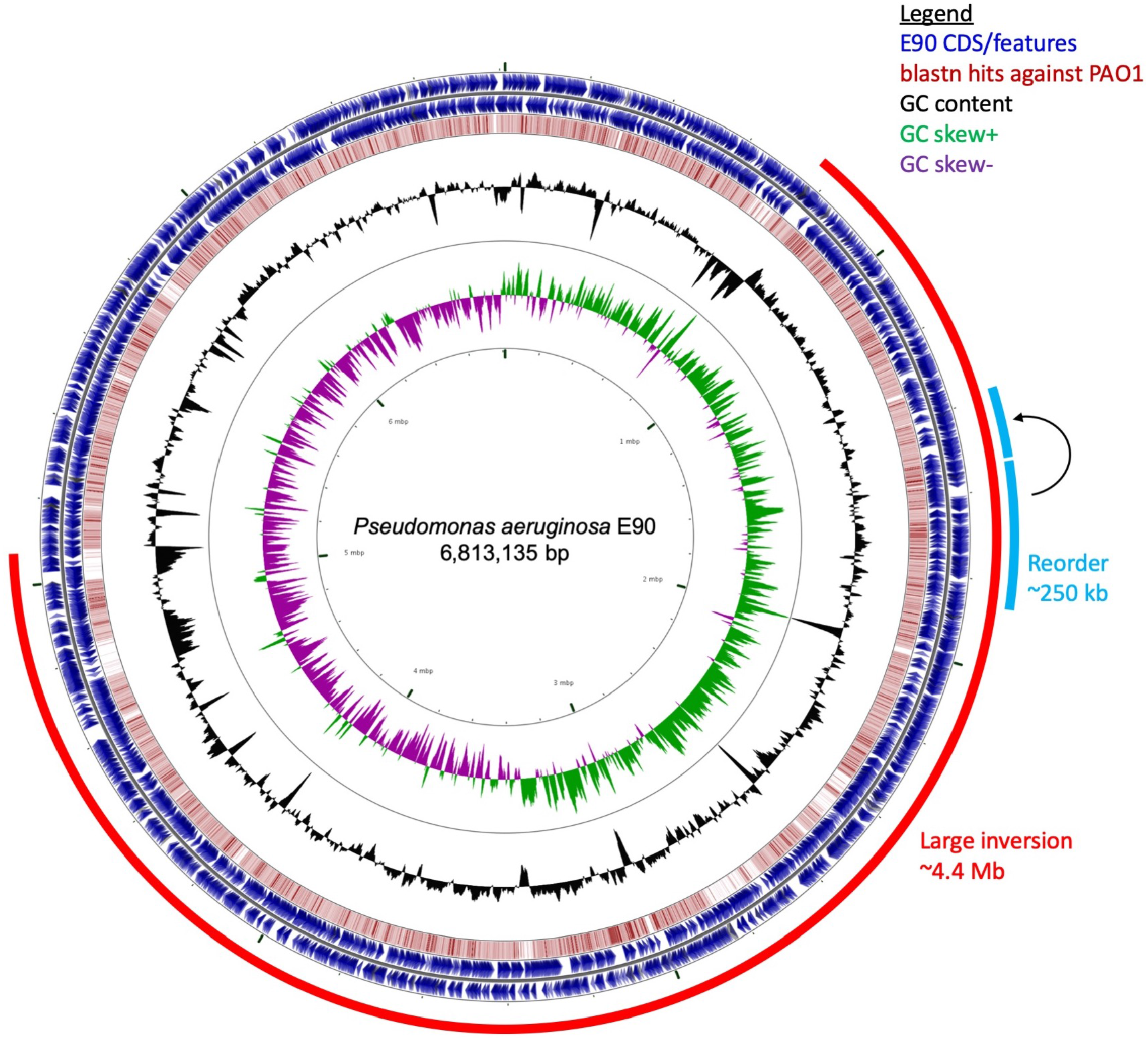
General features of the complete E90 genome. This circular representation of the E90 genome includes rings indicating the following features described from the outer-most to inner-most rings: annotated features of CDS (blue) or rDNA (grey) on the forward (outer) or reverse (inner) strands; all-by-all blastn hits (red) in a comparison against PAO1_107 (nucleotide identity >40%); GC content deviation (black); GC skew (+, green; -, purple). Additional outer, partial rings indicate the 4.4 Mb inversion (bright red) and the 250 kb reorder (light blue).

Next, to facilitate a comparison to previously published QS regulons in our transcriptome analysis [3,28], we grew strains in LB-MOPS to an OD_600_ of 2.0. The growth of E90 and E90Δ*rhlR* are indistinguishable in this medium (S1 Fig.). Our DE analysis identified 53 genes that were upregulated in the E90 vs. E90Δ*rhlR* comparison (Table 1 and S1 Table). Forty-four (83%) of these genes were identified as QS-regulated in a previous microarray study of the PAO1 [3] and 21 belong to the core quorum-controlled genome characterized in the laboratory strain PAO1 [28].

**Table 1.**
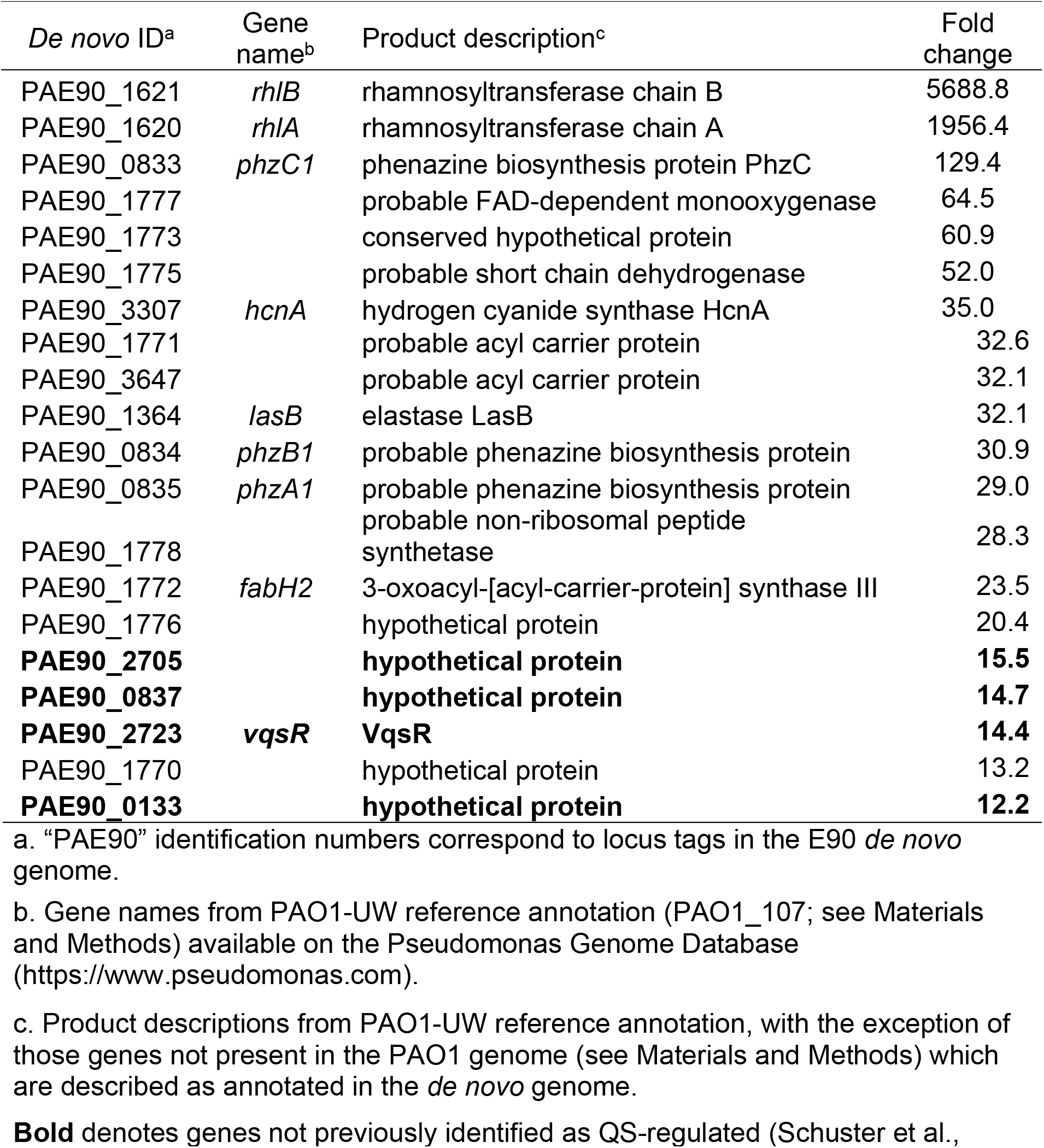
The 20 most highly RhlR-activated genes in isolate E90.

We also identified several well-known virulence genes including those that encode biosynthetic machinery required for rhamnolipid (*rhlAB*), hydrogen cyanide (HCN; *hcnABC*), elastase (*lasB*), and pyocyanin synthesis (*phzABC1*). Elastase is an exoprotease known to degrade various components of the innate and adaptive immune system including surfactant proteins A and D [29,30]. Rhamnolipid and pyocyanin have also been previously appreciated for their roles in airway epithelium infiltration and damage [31,32]. In addition, our RNA-seq analysis revealed *hsiA2*, the first gene in the cluster encoding the Second Type VI Secretion System, which facilitates the uptake of *P. aeruginosa* by lung epithelial cells [33].

While QS control of the phenazine biosynthesis pathway has been reported previously, only one of the two “redundant” operons (*“phz1”; phzA1-G1*) was indicated [3]. Interestingly, our transcriptome analysis found that RhlR also regulates the first two genes of the second phenazine operon (*“phz2”; phzA2-G2*) in E90, albeit at a slightly lower level than *phz1*. Both operons encode nearly identical sets of proteins, each with the capacity to synthesize the precursor (phenazine-1-carboxylic acid) of many downstream phenazine derivatives, including the virulence factor pyocyanin [34]. Despite their seemingly redundant function, *phz1* and *phz2* do not appear to be regulated in concert. In strain PA14, although *phz1* is more highly expressed than *phz2* in liquid culture, similar to what we observed in the E90 RhlR regulon, *phz2* actually contributes more to overall phenazine production in liquid culture [35]. Furthermore, *phz2* is the only active *phz* operon in colony biofilms, and was the only *phz* operon implicated in lung colonization in a murine model of infection [35].

Moreover, we observed that the RhlR regulon included genes that likely confer a growth advantage in the CF lung. For example, *cbpD* encodes a chitin-binding protein shown to contribute to the thickness of biofilms, the development of which is important for nutrient acquisition and stress resistance [36]. The gene encoding the monodechloroaminopyrrolnitrin 3-halogenase PrnC was also present in the E90 RhlR regulon, which has not been reported in previous *P. aeruginosa* transcriptomes and is not present in the PAO1 reference genome. Halogenase PrnC has only previously been described in *P. protegens* (formerly *P. fluorescens*), where it is involved in the synthesis of pyrrolnitrin, an antifungal antibiotic [37].

Among the most highly regulated genes [3,28] were those belonging to a conserved nonribosomal peptide synthetase (NRPS) pathway (PaE90_1770-1779; PA3327-3336). The products of this NRPS pathway have been identified as azetidine-containing alkaloids referred to as azetidomonamides[38]. The biological significance of this widely conserved NRPS pathway in *Pseudomonas* species or what roles azetidomonamides may play in virulence or interspecies interaction is not well understood, but regulation by QS appears to be a common feature.

Our interrogation of the E90 RhlR regulon also revealed 30 genes that were RhlR-repressed (S2 Table); none of these genes were reported in previous reports of QS-repressed genes [3] and 19 are not present in the PAO1 genome. We found two genes of the *alpBCDE* lysis cassette, *alpB* and *alpC*, were repressed by RhlR in E90 under the conditions of our experiments. While induction of *alpBCDE*, via de-repression of the *alpA* gene, has been shown to be lethal to individual cells, it may benefit infecting cells at the population level [39]. We also observed down-regulation of the gene encoding the posttranscriptional regulatory protein RsmA by RhlR in E90. RsmA is nested in a host of regulatory machinery important in infection, and mutation of RsmA has been observed to favor chronic persistence and increased inflammation in a murine model of lung infection [40]. Lastly, we identified RhlR regulation of phage loci not found in the PAO1 genome. The RhlR-repressed phage loci correspond to E90 genes PaE90_2433 through PaE90_2442.

### RhlR is the primary driver of cytotoxicity in a lung epithelium model

LasR-null laboratory strains are less virulent than the WT in acute infection settings [10,11,41]. However, as the RhlR-dependent QS regulon of E90 includes several factors implicated in virulence (Table 1), we queried if E90 might be capable of inducing host cell death. To address this question, we incubated an *in vivo*-like three-dimensional (3D) lung epithelial cell culture model (A549 cell line) [42] with either PAO1, E90, or engineered QS transcription-factor mutants. The 3D lung cell model possesses several advantages over the standard A549 monolayer as an infection model, including increased production of mucins, formation of tight junctions and polarity, decreased expression of carcinoma markers, and physiologically relevant cytokine expression and association of *P. aeruginosa* with the epithelial cells [42,43]. Following an incubation period of 24 hours, we measured cell death of the 3-D cell cultures via cytosolic lactate dehydrogenase (LDH) release. Consistent with prior studies, WT PAO1 cytotoxicity is abrogated in a LasR deletion mutant; however, cytotoxicity of a PAO1 RhlR-null mutant is similar to that of the wild-type, because in this assay the secreted products responsible for cytotoxicity are LasR-regulated in PAO1, with little or no contribution from RhlR. Strikingly, the opposite was true for E90: deletion of RhlR significantly reduces cytotoxicity (Fig. 5). This RhlR-dependent cytotoxicity might be related to the different timing of RhlR activation in E90, the specific set of genes regulated by RhlR in this strain, or both. Together, these results highlight the restructuring of QS gene regulation in this clinical isolate and underscore implications for virulence during chronic infection.

**Figure 5.**
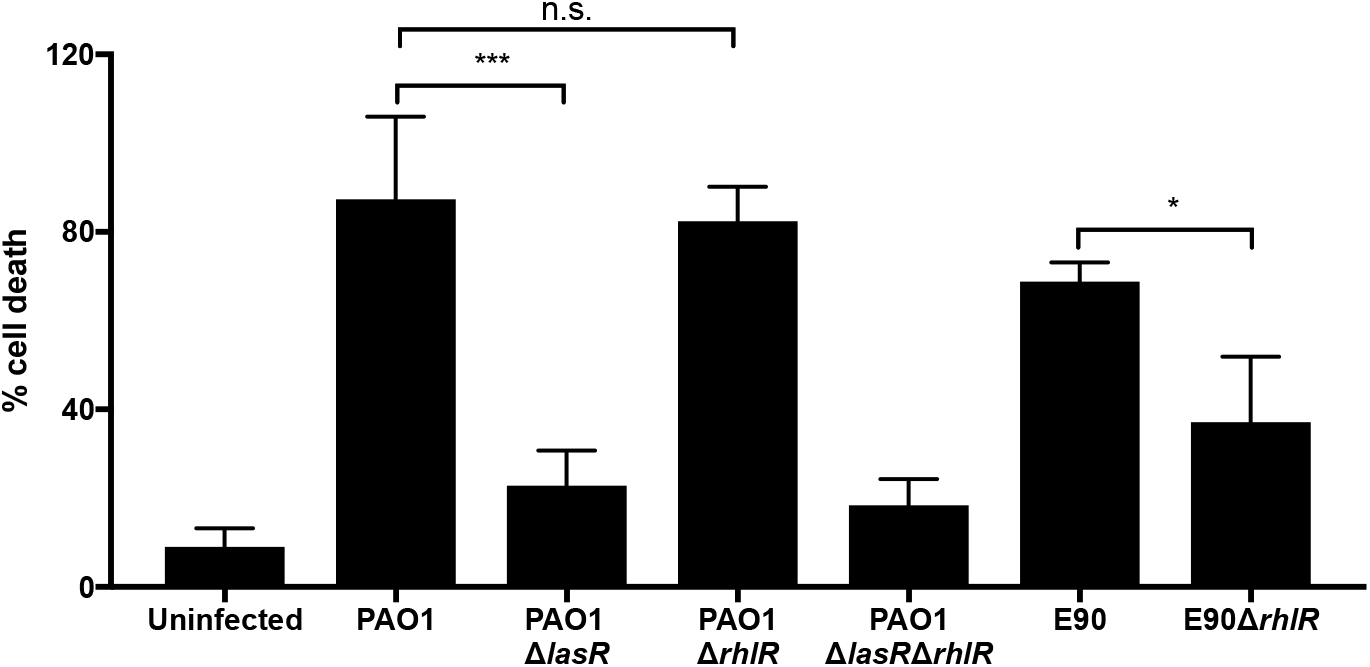
RhlR regulates cytotoxicity in E90, but not PAO1. We measured cell lysis (as a percentage of the total lactate dehydrogenase release caused by incubation with a lysis agent) of A549 cells incubated with PAO1, E90, or QS transcription factor mutants of PAO1 or E90. Asterisks (*) indicate statistical significance (*, p < 0.05; **, p < 0.01; ***, p < 0.001 by t-test). Error bars represent the standard deviation for results of at least three independent experiments.

## DISCUSSION

A substantial body of literature now suggests that the QS hierarchy of *P. aeruginosa* is adaptable and that LasR mutants can be “rewired” to be AHL QS proficient [16,25,26,44]. These “rewired” LasR-null clinical isolates retain the QS regulation of several exoproducts through the RhlI/RhlR circuit [16,17]. Prior studies examining a RhlR-dependent variant of PAO1 [24], and another LasR-null and RhlR-active clinical isolate[17], show that the parent strain outcompetes RhlR-null derivatives when grown in co-culture [17,24]. These findings support the notion that there is something inherently disadvantageous about mutation of RhlR and point to RhlR as a key QS transcription factor in chronic infections like CF.

We do not know the mechanism or genetic modifications that resulted in Las-independent RhlR activity in isolate E90. In strain PAO1, in which the hierarchy of QS was initially described, LasR mutants can readily evolve an independent RhlR QS system through inactivating mutations of *mexT*, which encodes a non-QS transcriptional regulator [24,26]. However, this is not the case in isolate E90, which possesses a functional *mexT* allele. We did observe that *rhlI* expression is upregulated by RhlR in E90 unlike in PAO1, where *rhlI* expression is predominately LasR-regulated [45]. These data suggest that in E90 RhlR and RhlI may constitute a positive autoregulatory loop that may facilitate Las-independent RhlR activity. We are interested in investigating alternate mechanisms, other than inactivation of *mexT*, through which RhlR escapes LasR regulation in these “rewired” backgrounds.

In the present study, we aimed to identify which genes comprise the RhlR regulon in a clinical isolate, which may shed light on factors important for establishment or continuation of a chronic infection. Our RNA-seq analysis revealed that the E90 RhlR regulon bears a substantial amount of overlap with the suite of AHL-regulated genes previously identified in PAO1[3] and consists of virulence factors that are likely advantageous in the context of the CF lung.

A portion of the genes found to be RhlR-regulated in our transcriptomic analysis of E90 were not previously reported to be QS-regulated. These genes include *vqsR* (PAE90_2723/PA2591) and two genes of the *phz2* operon (PAE90_3614-3615/PA1899-1900). VqsR is itself a LuxR-homolog that serves to augment QS gene regulation, possibly through activation of the orphan QS receptor QscR, although the precise mechanism and biological outcomes of this interaction are still mysterious [46,47]. Our finding that the *phz2* operon, in addition to *phz1*, is activated by RhlR may reflect ongoing QS adaptation in our selected CF isolate. While E90 appears to produce slightly less bulk pyocyanin in broth culture than PAO1, pyocyanin production by E90 may be comparatively greater in the biofilm lifestyle of the CF lung. The *phz2* locus, while showing roughly 98% nucleotide identity with *phz1*, has been shown to be responsible for nearly all the pyocyanin produced in biofilms by PAO1 and is the dominant contributor to murine lung colonization between the two loci [35]. It is possible that some of these previously unreported QS-regulated genes were excluded from earlier transcriptome analyses[3,28] due to different analysis approaches or methodology. Of particular note, we compared a RhlR-deletion mutant to the parent strain to derive our transcriptome, while some of these previous studies used signal-synthase mutants with and without signal, which has been demonstrated, in the case of RhlR QS, to yield a different phenotype [48].

We also discovered RhlR-QS-regulation of many genes that are not present in the PAO1 genome. This list includes several hypothetical proteins activated as much as 15-fold in E90 compared to the RhlR mutant. The list also includes the gene encoding the halogenase PrnC, a protein involved in production of the antifungal antibiotic pyrrolnitrin which may be important in interspecific interactions in the CF lung [49]. Our finding that RhlR-QS in E90 also appears to repress genes in the programmed cell death cassette *alpBCDE*, points to additional potential for QS regulation of population level interactions in CF-adapted strain E90.

Although we do not yet fully understand the biological significance of the RhlR-mediated suppression of the phage identified in this study, we are interested in exploring its role, if any, in fitness and inter- and intra-species competition in the near future. We note that had we used the PAO1 genome, as opposed to the E90 *de novo* genome, for read alignment, we would have failed to identify the phage loci and a handful of other genes. These findings therefore argue in favor of using *de novo* genomes to improve comprehensive transcriptome analyses of clinical and environmental isolates moving forward.

Strikingly, we found that in E90, RhlR but not LasR is the critical determinant of cytotoxicity in a three-dimensional lung epithelium cell aggregate model. Though our study did not reveal exactly which virulence factors are important for cell death in this model, our results nevertheless challenge the idea that LasR-null isolates are avirulent. Instead, our data argue that some virulence activity is conserved in “rewired” isolates, but that RhlR is the primary regulator of several such functions.

The scope of our analysis is limited by our examination of a single clinical isolate and laboratory growth conditions were used for RNA-seq analysis, but our data provide a basis for understanding regulatory remodeling of QS activity and provide avenues for future investigation. Several important questions remain about QS in clinical isolates, including whether or not there is a “core” regulon that is common to isolates that use either LasR or RhlR as the primary QS transcription factor. Our work also serves as a starting point to test hypotheses regarding the role of RhlR-regulated genes during chronic infection, the possible fitness advantage associated with LasR-independent RhlR activity, and mediators of sustainable chronic infections. Our work reveals the potential breadth of QS activity and virulence functions retained in LasR-null, CF-adapted isolates and suggests that the development of anti-QS therapeutics for chronic *P. aeruginosa* infections should be focused on RhlR, not LasR.

## MATERIALS AND METHODS

### Bacterial strains and growth conditions

Bacterial strains and plasmids used in this study are described in S3 Table in Supporting Information. E90 is part of a collection of clinical isolates obtained via the Early *Pseudomonas* Infection Control Observational (EPIC Obs) Study[20]. The isolates are from oropharyngeal and sputum samples from 5-12 year-old patients. Further details regarding the EPIC Obs study design and results have been described previously [20,50].

For the transcriptional reporter assays as well as pyocyanin and AHL measurements, overnight cultures were started from single colonies grown in 3 mL of Luria-Bertani (LB) broth buffered with 50 mM morpholinopropanesulfonic acid (MOPS) in an 18 mm culture tube. For the cytotoxicity experiments, overnight cultures were started from single colonies in 5 mL of unbuffered LB broth. When appropriate, antibiotics were added at the following concentrations: 10 μg/mL gentamicin or 100 μg/mL ampicillin for *Escherichia coli*, and 100 μg/mL gentamicin for *P. aeruginosa*. Cells were grown at 37°C with shaking at 250 RPM unless stated otherwise.

### LasR and RhlR activity

LasR and RhlR-specific promoter fusions constructed in pPROBE-GT have been described previously [16] and are listed in S3 Table. Electrocompetent *P. aeruginosa* cells were prepared through repeated washing and resuspension of cell pellets in 300 mM sucrose[51]. Transformants were obtained by plating on LB agar supplemented with gentamicin and verified by PCR.

Experimental cultures were prepared as follows: first, overnight cultures were grown with the addition of 100 μg/mL gentamicin and 100 μg/mL AiiA lactonase, the latter inhibiting AHL-mediated QS[22]. The addition of AiiA lactonase eliminates residual GFP fluorescence that would otherwise arise from previously induced reporter gene expression during overnight growth. Overnight cultures were then diluted to an optical density (OD_600_, 1 cm pathlength) of 0.001 (approximately 1-5×10^6^ CFU/mL) in 3 mL MOPS-buffered LB supplemented with AiiA lactonase in 18 mm culture tubes. After these cultures grew to an approximate OD_600_ of 0.2, they were diluted to OD_600_ 0.001 in 400 μL of MOPS-buffered LB alone in a 48-well plate with a clear bottom (Greiner Bio-One). To prevent evaporation, strains were only grown in wells that did not line the edges of the plate and all empty wells were filled with 400 μL water. We monitored GFP fluorescence and OD_600_ at 30-minute intervals for 15 hours using a BioTek Synergy HI microplate reader (excitation: 489 nm, emission: 520 nm, gain: 80). All strains were grown at 37°C with shaking for the duration of the assay. To account for differences in growth, results were normalized to OD_600_. As a negative control, each strain was electroporated with an empty vector, which was used to establish a baseline level of background fluorescence. The fluorescence intensity was calculated by subtracting the background fluorescence from the total fluorescence measured at every time point. All experiments were performed in biological triplicate.

### Construction of the E90Δ*rhlR* mutant

A homologous recombination approach was used to generate an in-frame deletion mutant[52,53]. Fragments flanking *rhlR* were PCR-amplified from E90 genomic DNA and cloned into pEXG2 to yield pEXG2.E90Δ*rhlR*, which was then transformed into *E. coli* S17-1 in order to facilitate conjugal transfer of pEXG2.E90Δ*rhlR* into E90. Transconjugants were selected by plating on *Pseudomonas* isolation agar supplemented with gentamicin, and deletion mutants were counter-selected by plating onto LB agar with 10% (wt/vol) sucrose. Deletion of *rhlR* was confirmed by PCR and targeted sequencing.

### AHL signal extraction and measurement

Experimental cultures were prepared from overnight cultures diluted to OD_600_ 0.001 in 3 mL of MOPS-buffered LB in an 18 mm culture tube. Experimental cultures were grown with shaking until they reached OD_600_ 2.0. Then, AHL signals were extracted from experimental cultures using acidified ethyl acetate as described elsewhere[54]. We used an *E. coli* DH5α strain containing either pJN105L and pSC11 in conjunction with the Tropix^®^ Galacto-Light™ chemiluminescent assay (Applied Biosystems) to measure 3OC12-HSL, or containing pECP65.1 to measure C4-HSL[23,55,56]. The bioassay strains and plasmids are listed in S3 Table in the Supporting Information.

### Protease and pyocyanin measurements

Experimental cultures were prepared from overnight cultures by diluting to OD_600_ 0.001 in 3 mL MOPS-buffered LB in 18 mm culture tubes. For secreted protease measurements, experimental cultures were grown with shaking to OD_600_ 2.0. Then, cells were pelleted and 100 μL of the supernatant was collected to measure protease production using the FITC-Casein for Pierce Fluorescent Protease Assay Kit (ThermoFisher Scientific). For pyocyanin measurements, experimental cultures were grown with shaking for 18 h and pyocyanin was extracted from cultures as described previously[16]. We grew strains in MOPS-buffered LB to remain consistent with the growth conditions used for RNA-seq analysis.

### Cytotoxicity of three-dimensional A549 cell cultures

A three-dimensional lung epithelial cell culture model was generated by culturing A549 cells (ATCC CCL-185) on porous microcarrier beads in a rotating well vessel (RWV) bioreactor system, as described previously [42]. A549 cells were grown in GTSF-2 medium (GE Healthcare) supplemented with 2.5 mg/L insulin transferrin selenite (ITS) (Sigma-Aldrich), 1.5 g/L sodium bicarbonate, and 10% heat-inactivated FBS (Invitrogen), and incubated at 37°C under 5% CO_2_, >80% humidity conditions. Infection studies were performed on cultures grown for 11 to 14 days in the RWV. Thereafter, 3-D cell cultures were equally distributed in a 48-well plate at a concentration of 2.5 × 10^5^ cells/well (250 μL volume), and infected with the different strains at a targeted multiplicity of infection of 30:1 as described previously[43]. All infection studies were performed in the above-described cell culture medium, with the exception that no FBS was added given the interference of serum compounds with QS signaling [57]. After 24 h infection, the release of cytosolic lactate dehydrogenase (LDH) from 3-D lung epithelial cell cultures was determined using a LDH activity assay kit (Sigma-Aldrich) according to the manufacturer’s instructions. A standard curve using NADH was included. The positive control (theoretical 100% LDH release) was obtained by lysing 2.5 × 10^5^ cells with 1% Triton-X100. All LDH release values were expressed as a percentage of the positive control.

### RNA isolation and qRT-PCR

Overnight cultures were started from single colonies grown in 3 mL of MOPS-buffered LB in 18 mm culture tubes. Experimental cultures were prepared by diluting overnight cultures to OD_600_ 0.01 in 25 mL of MOPS-buffered LB in 125-mL baffled flasks. Experimental cultures were grown at 37°C with shaking at 250 RPM. Approximately 1 x 10^9^ cells were pelleted at OD_600_ 2.0 and mixed with RNA Protect Bacteria reagent (Qiagen) before being stored at −80°C. Thawed cell pellets were resuspended in QIAzol reagent and mechanically lysed by bead beating. To extract RNA, we used the RNeasy kit (Qiagen) according to manufacturer’s instructions. Isolated RNA was then treated with Turbo DNAse (Ambion) and purified using the MinElute cleanup kit (Qiagen). Three biological replicates were processed for each strain (E90 and E90Δ*rhlR*). Next, cDNA was prepared using the iScript™ cDNA Synthesis Kit (BioRad). Then, expression of target genes was analyzed by following the protocol for the iQ™ SYBR^®^ Green SuperMix (BioRad) on a CFX96 Real-Time PCR cycler. We analyzed expression of the following genes: *lasI, lasB, rhlI, rhlR, rhlA, pqsA, chiC*, and *aprA*. We used *rplU* as a reference gene. Primers used for qRT-PCR are listed in S3 Table in Supporting Information.

### Whole-genome sequencing, RNA-seq, and differential gene expression analysis

We generated the complete circular sequence of E90 using a *de novo* whole-genome sequencing approach. High-molecular weight (HMW) genomic DNA was isolated from overnight E90 liquid culture using the Genomic-tip 20/G kit (Qiagen). Genomic DNA was sequenced separately using the following two approaches. For short reads, genomic DNA was subjected to 300 bp PE sequencing on the Illumina MiSeq platform using TruSeq v3 reagents to yield approximately 20 M raw reads which were then groomed using Trimmomatic (v0.36; adapter trimming, paired reads only, Phred score cutoff=15) [58]. For long reads, genomic DNA was prepared into two ligation-mediated (SQK-LSK109, Oxford Nanopore) libraries: one barcoded via PCR (EXP-PBC001) and the other via native barcoding (EXP-NBD114). Libraries were then subjected to sequencing on the Nanopore MinION platform using R9.4.1 pores. Nanopore reads were base-called and de-multiplexed using Guppy (v3.1.5, Oxford Nanopore), further groomed to remove adapters and for quality using Porechop (v0.2.4) [59], and final statistics determined in NanoPack (NanoPlot v1.27.0; NanoQC v0.9.1) [60] (read length N50=12kb; median read quality=Q12.6). All reads were then combined in a hybrid *de novo* assembly approach using the Unicycler pipeline [61], including short-read assembly via SPAdes(v3.13.0)[62], long-read assembly via Racon (v1.4.3), and polishing via Pilon [63], to yield the complete E90 genome. The E90 genome was then annotated using the RAST pipeline [64]. The E90 genome is available via the National Center for Biotechnological Information (NCBI) under BioProject accession PRJNA559863.

For RNA-seq experiments, cultures were prepared, and RNA was extracted and purified as described above for qRT-PCR with 2 biological replicates per treatment. Genewiz, LLC performed rRNA depletion, library generation, and sequencing for all samples. RNA reads were obtained using the Illumina HiSeq platform with an average of 15.3M 150-bp paired-end raw reads per sample which were then groomed using Trim Galore (v0.4.3; https://github.com/FelixKrueger/TrimGalore). Reads were then aligned against the E90 genome and counted using the Subread/featureCounts suite of command line tools to produce a final count matrix of 4 by 6478 which was then loaded into the R statistical environment [65,66]. Differential expression (DE) analysis was performed using DESeq2 using a fold-change cut-off of 2 and an adjusted p=0.05 [67]. The raw reads and count matrix associated with this transcriptome analysis have been deposited in the of the NCBI Sequence Read Archive under BioProject accession PRJNA559863.

## ACKNOWLEDGMENTS

This work was supported by grants from the NIH (R01 GM125714), Doris Duke Charitable Foundation (2017072) and the Burroughs-Wellcome Fund (1012253) to AAD. RC and KLA were supported in part by the Cystic Fibrosis Foundation, with additional support to KLA from the US National Institutes of Health (T32 HL007287). We acknowledge core support from the Cystic Fibrosis Foundation (SINGH15R0 and R565 CR11) and NIH (P30DK089507). Funding from the Research Foundation Flanders to AC (Odysseus grant G.0.E53.14N) and SVDB (PhD fellowship 3S55719) also supported this study. We thank Amy Schaefer and Nicole Smalley for providing the AiiA lactonase and protocols for the use of AiiA lactonase.

## SUPPORTING INFORMATION

**S1 Fig. Growth curves of E90 and E90Δ*rhlR* in buffered Luria-Bertani Broth.** Means and standard deviation of biological replicates are shown (n=3). In some cases, error bars are too small to be seen.

**S1 Table. Complete list of RhlR-activated genes in strain E90.**

**S2 Table. Complete list of RhlR-repressed genes in strain E90.**

**S3 Table. Bacterial strains, plasmids, and primers used in this study.**

